# Finding the Goldilocks zone: Modulating glycoprotein cleavage and fusogenicity optimizes the efficacy of a candidate Crimean-Congo hemorrhagic fever virus vaccine

**DOI:** 10.64898/2026.06.07.730762

**Authors:** Jacob Berrigan, Stephanie R. Monticelli, Pieter Spealman, Courtney A. Cohen, Thomas G. Batchelor, Albert Wang, Elizabeth McFadden, Russell R. Bakken, Megan M. Slough, Ariel S. Wirchnianski, Eva Mittler, Robert W. Cross, Thomas W. Geisbert, Jesse H. Erasmus, David W. Hawman, Jason S. McLellan, Andrew S. Herbert, Kartik Chandran

**Author notes:** Equal contributors (Jacob Berrigan, Stephanie R. Monticelli). Corresponding authors: K.C., A.S.H.

## Abstract

Crimean-Congo hemorrhagic fever virus (CCHFV) is the etiologic agent of a lethal hemorrhagic disease spread by ticks throughout Europe, the Middle East, Africa and Asia. The lack of approved medical countermeasures and fundamental understanding of molecular mechanisms of viral assembly and egress have thus far curtailed disease prevention. Here, we identify and characterize key residues within the viral glycoprotein through forward and reverse genetics for vesicular stomatitis virus (VSV)-based vaccine candidates that are highly protective in animal models. These residues are broadly applicable across divergent CCHFV strains and lead to greater protection *in vivo* against heterologous challenge. We further characterize the essential role of proteolytic processing in the maintenance of a stable fusogenic state required for effective VSV-based CCHFV vaccines. This study establishes a toolkit for better understanding orthonairovirus glycoprotein processing and vaccine development.

## Introduction

Crimean-Congo hemorrhagic fever virus (CCHFV) is a member of the family *Nairoviridae* in the class *Bunyaviricetes* of enveloped, segmented, negative-strand RNA viruses. Carried mainly by hyalommid ticks, CCHFV is the most prevalent tick-borne virus that causes human disease and is endemic in countries across Europe, the Middle East, southern Asia, and Africa [1–3]. The extraordinary geographic diversity of CCHFV is matched by its sequence diversity—CCHFV isolates have been grouped into seven distinct genetic clades (Africa 1, Africa 2, Africa 3, Asia 1, Asia 2, Europe 1 and Europe 2) [4] (S1 Fig). CCHFV is transmitted to humans by tick bite or contact with infected blood or tissues. In endemic regions, this often occurs through contact with domesticated ungulates such as goats and cattle infected with CCHFV. Case-fatality rates of 5–30% are typically observed in CCHF outbreaks but have been reported to be as high as 80% [2, 5, 6]. Disease onset is rapid and initially characterized by high fever, joint pain and vomiting, followed by severe bruising and uncontrollable bleeding[1]. Leukocytosis, abdominal pain, and diarrhea are common in fatal cases. No FDA- or EMA-approved specific medical countermeasures are currently available for CCHF. The broad-spectrum antiviral ribavirin is used off-label in some regions, but evidence for its efficacy is controversial [7]. In 2017, the World Health Organization declared CCHF a Blueprint Priority Disease [8].

The enveloped virions of CCHFV package a tri-segmented negative-strand RNA genome. The medium (M) segment encodes a large membrane-embedded polyprotein, pre-GPC, which is a major target of the antiviral immune response [9]. Pre-GPC undergoes maturation through a complex cascade of proteolytic processing events. It is co-translationally cleaved by host signal peptidase to pre-Gn, NSm, and pre-Gc in the endoplasmic reticulum (ER) [10, 11]. Pre-Gn:pre-Gc complexes are subsequently trafficked to the Golgi apparatus where they are further cleaved by the proprotein convertases subtilisin kexin isozyme-1/site-1 peptidase (SKI-1/S1P; hereafter, SKI-1) and furin. Sequential SKI-1 and furin cleavages of pre-Gn yield the N–terminal mucin-like domain, GP38, and Gn [12–15]. A currently unknown proprotein convertase cleaves pre-Gc to Gc [11]. We recently showed that the pre-fusion conformer of the entry glycoprotein complex consists of a ternary complex of Gn, Gc, and GP38 [16]; however, the higher-order organization of this complex at the virion surface remains unknown, as are the precise structural rearrangements in this complex during virion biogenesis and cell entry. A non-virion–associated soluble form of GP38 is also produced by infected cells [10]. Whether this soluble GP38 is derived from immature GPC complexes or liberated from mature, virion-associated glycoprotein complexes, or both, is unclear at present. We recently uncovered a novel non-entry–related role for GP38 as a viral toxin that destabilizes endothelial barrier function, causing endothelial leak [17].

Although no FDA- or EMA-approved vaccines are currently available for CCHFV, efforts to develop such vaccines go back decades. An early live-attenuated vaccine, generated by viral serial passage in the brains of suckling mice, is only approved in Bulgaria to immunize military and medical personnel [18]. This vaccine has significant drawbacks, including the need for a four-dose series and boosts every five years. More recent work has leveraged the immunogenicity of GPC and the viral nucleoprotein (NP) across a variety of vaccine platforms, including recombinant vesicular stomatitis (rVSV) and adenovirus vectors, virus-like particles (VLPs), protein subunits, DNA, mRNA, and self-amplifying RNA [16, 19–24].

Only the rVSV and VLP platforms have been demonstrated to provide complete protection with a single dose of a GPC-based immunogen [19, 21, 25]. The capacity of rVSVs to replicate *in vivo*, unlike VLPs, makes rVSVs especially promising vaccine candidates. Rodriguez et al. showed that an rVSV vector bearing a VSV-adapted GPC as its only entry glycoprotein could elicit neutralizing antibodies and fully protect immunodeficient mice against lethal CCHFV challenge Independently, Tipih et al. (2025) also showed that an rVSV-vector bearing an overlapping but distinct suite of amino acid substitutions in GPC conferred partial protection (at a lower vaccine dose) in a transiently immunosuppressed murine model [19, 25]. The molecular and cellular basis of GPC adaptation to a replicating VSV remains poorly understood, as are the molecular mechanisms by which these adaptations afford enhanced GPC immunogenicity and vaccine-mediated protection.

Herein, we investigated these mechanisms through a systematic dissection of the rVSV-CCHFV vector described by Rodriguez and co-workers [19]. Using long-read sequencing, we uncovered two additional GPC amino acid substitutions not identified in the original report. Using VSV forward genetics, we defined a minimal suite of four essential GPC substitutions and demonstrated that it is necessary and sufficient for VSV adaptation of a divergent CCHFV GPC. This new VSV vector conferred protection against heterologous challenge and elicited high levels of neutralizing and non-neutralizing antibodies targeting Gc and GP38, respectively. Our mechanistic studies provide the first evidence that SKI-1 cleavage at the GP38-Gn boundary modulates the pH threshold of CCHFV membrane fusion and point to a key role for virion-associated GP38 in regulating the pre-fusion conformation of the viral glycoprotein spike during glycoprotein biogenesis, viral egress, and cell entry.

## Results

### Generation of a replication-competent vesicular stomatitis virus bearing CCHFV GPC and a fluorescent reporter

Previous work by Rodriguez and co-workers had described a replication-competent vesicular stomatitis virus (rVSV) encoding CCHFV GPC replacing the native G protein, generated by reverse genetics and serial passage [19]. Because this virus lacked a reporter, we sought to facilitate the study of rVSV-CCHFV assembly and entry by incorporating a mNeongreen-VSV phosphoprotein gene fusion (mNg-P) in the viral genome, as described [26] (Fig 1A). Accordingly, we engineered a VSV genome construct bearing mNg-P and a human codon-optimized IbAr10200 (hereafter, IbAr) GPC sequence encoding the putative adaptive changes (S368T, L518V, L1638R, R1648Q, and a 14-amino acid deletion at the C-terminus) described previously [19]. Unexpectedly, however, we were unable to recover an rVSV encoding this GPC construct using plasmid-based reverse genetics. To account for the possibility that one or more essential changes in the published vector might have been missed, we amplified the GPC gene by reverse transcription-PCR (RT-PCR) from a stock and introduced this sequence into our viral construct [19]. Strikingly, we were immediately successful in rescuing this virus (hereafter, rVSV-CCHFV GPC) (Fig 1B). Plaque-purified isolates of rVSV-CCHFV GPC efficiently infected Vero cells and produced high titers of infectious virus in cell supernatants (2±1×10^7^ infectious units (IU) per mL). Large-scale preparations of rVSV-CCHFV GPC were generated by infecting Vero cells at a multiplicity of infection (MOI) of 0.01 infectious units (IU)/cell for 2 days. rVSV-CCHFV GPC was neutralized by three previously described Gc-specific human monoclonal antibodies (mAbs) [27–29] (Fig 1C), providing evidence that infection by this surrogate viral vector reflects the authentic cell entry activity of CCHFV GPC.

**Figure 1.**
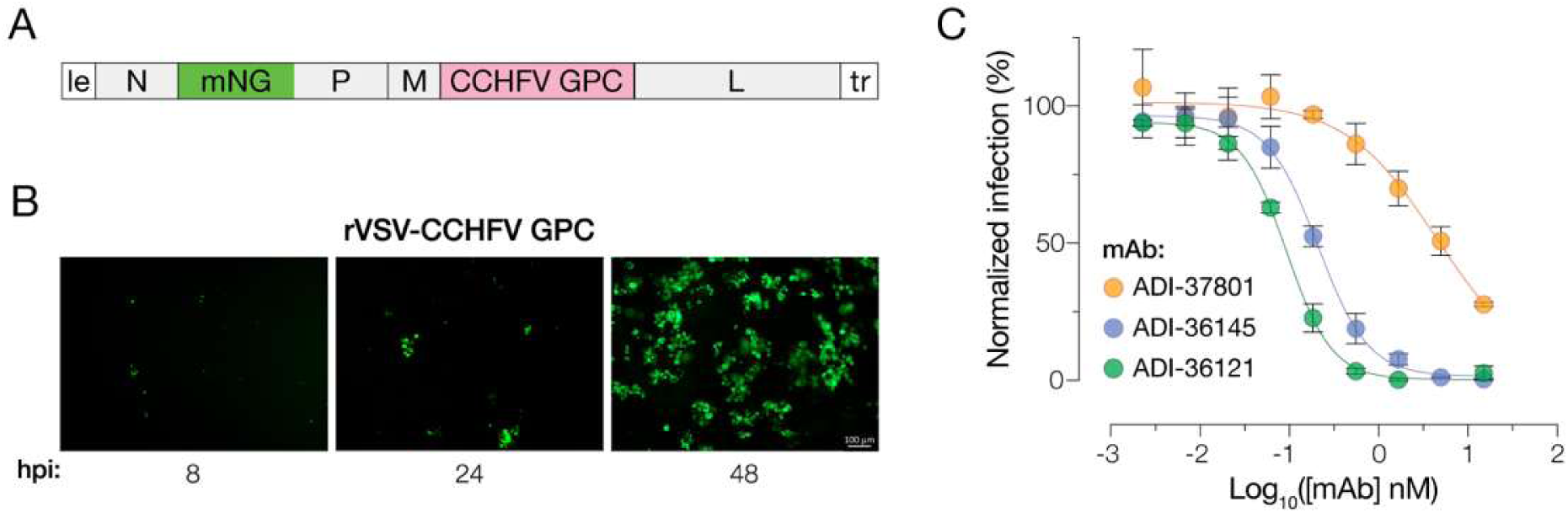
Rescue and characterization of a recombinant VSV bearing CCHFV GPC (rVSV-CCHFV GPC). **(A)** Organization of the rVSV-CCHFV GPC (IbAr10200) genome. Le, leader. N, nucleoprotein. mNg-P, mNeongreen reporter fused to phosphoprotein. L, polymerase. tr, trailer. **(B)** HUH7.5.1 cells were exposed to rVSV-CCHFV GPC and imaged by fluorescence microscopy at the indicated h post-infection (hpi). Representative images are shown. **(C)** rVSV-CCHFV GPC was pre-incubated with the indicated monoclonal antibodies (mAbs) and then exposed to Vero cells. Infection was scored by mNg expression at 16–18 h post-infection. Means ± s.d. from two independent experiments are shown (*n*=6).

### rVSV-CCHFV GPC bears two additional amino substitutions in the Gn ectodomain

To test the hypothesis that rVSV-CCHFV GPC bears one or more essential amino acid substitutions not described previously, we sequenced an RT-PCR product of GPC derived from viral genomic RNA with the Oxford Nanopore (ONT) platform. We identified the four missense mutations and the deletion observed by Rodriguez et al. (2019). However, we also uncovered two additional missense mutations encoding amino acid substitutions D525G and Y728S in the Gn ectodomain and Gn cytoplasmic tail (C–tail), respectively (Fig 2, S1 Fig and S3 Fig).

**Figure 2.**
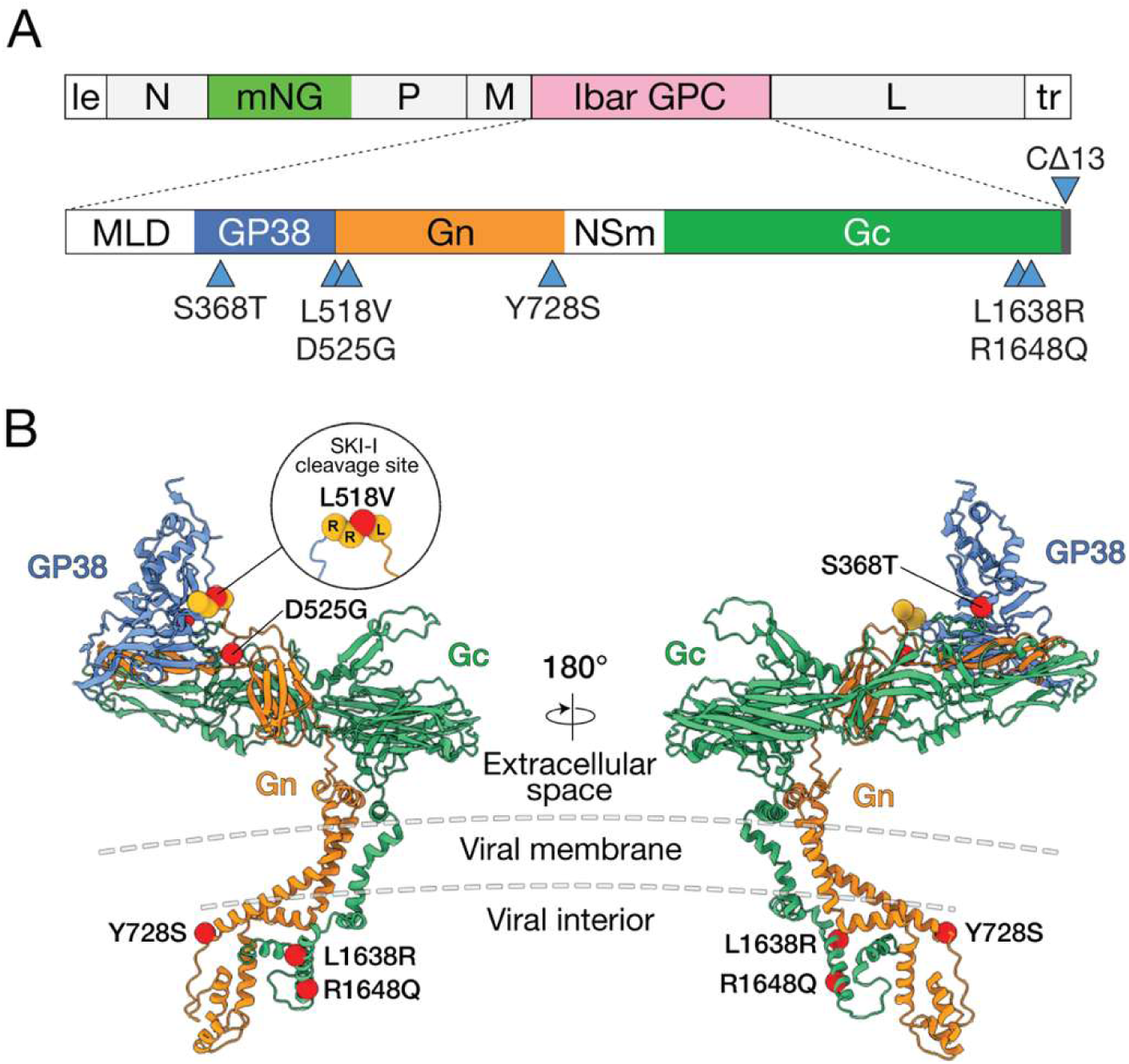
Identification of amino acid substitutions in rVSV-CCHFV GPC by long-read sequencing. **(A)** cDNA derived from a rVSV-CCHFV GPC population was subjected to long-read sequencing, genome assembly, and variant calling. Significantly enriched amino acid substitutions are shown. **(B)** Ribbon representation of CCHFV GPC pre-fusion structure generated by AlphaFold 3 with mutations in **(A)** highlighted.

### Identification of adaptive amino acid substitutions in rVSV-CCHFV GPC

We adopted a forward-genetic approach to assess the importance of each of the six amino acid substitutions in rVSV-CCHFV GPC for viral infection and multiplication. Specifically, we engineered a panel of rVSV-CCHFV GPC genome plasmids in which each amino acid was individually reverted to wild-type (WT) (Fig 3A). We then transfected these plasmids into cells along with helper plasmids to generate rVSVs [30]. Viral stocks were serially passaged three times in Huh7.5.1 cells. cDNAs generated from viral supernatants collected after each passage were barcoded and subjected to Oxford Nanopore long-read sequencing (Fig 3B). Sequence analysis revealed that L518V (SKI-1 cleavage site) and L1638R (Gc C–tail) re-emerged rapidly in the rescue population (Fig 3C), indicating that these substitutions are critical for rVSV-CCHFV GPC rescue. D525G (GP38-Gn interface) and Y728S (Gn C–tail) were enriched more gradually but were present in >80% of sequences by passage 3, providing evidence that these substitutions confer a significant fitness benefit. By contrast, the WT residues at positions 368 (GP38) and 1648 (Gc C–tail) were observed in the majority of the reads at passage 3, suggesting that S386T and R1648Q are dispensable for rVSV-CCHFV GPC rescue and multiplication and may be hitchhiker mutations with no meaningful phenotypic effect.

**Figure 3.**
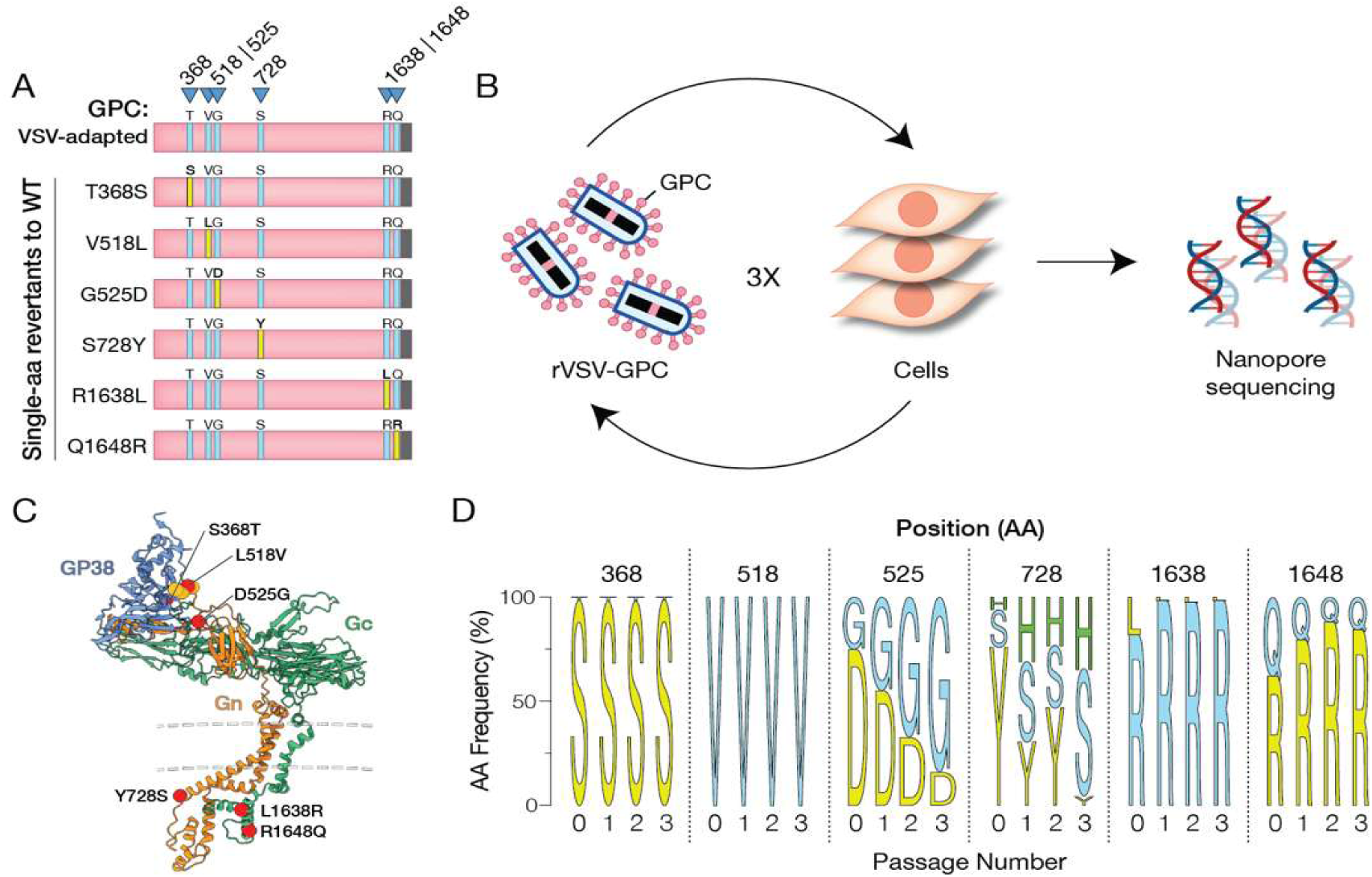
Forward-genetic studies to identify adaptive substitutions in rVSV-CCHFV GPC. **(A, C)** GPC constructs bearing six single-amino acid reversions to WT used for the forward genetic studies in **(B–D)** are shown. Yellow and blue rectangles indicate WT and mutant residues, respectively, at the indicated positions.**(B)** Schematic of experimental workflow for rVSV-CCHFV GPC adaptation by serial passage. rVSVs were rescued and subsequently subjected to blind serial passage in Vero cells three times. Progeny viral genomes were sequenced as in Fig 2A. **(D)** Amino acid residue frequencies at each position in the starting (passage 0) and serially passaged populations (passages 1–3) are represented as logo plots. The results of two independent experiments are shown.

### Substitutions D525G and/or Y728S enhance rVSV infectivity by increasing GPC incorporation into viral particles

To further dissect the fitness advantage afforded by the four key substitutions identified above, we sought to rescue rVSV-CCHFV GPCs bearing only the two essential substitutions (L518V and L1638R; 2-Mut) or bearing all four substitutions (4-Mut). Both sets of amino acid changes were introduced into GPC in the background of the C–terminal deletion (Fig 4A). We successfully recovered both rVSVs. However, rVSV-CCHFV GPC(4-Mut) infected cells much more efficiently than rVSV-CCHFV GPC(2-Mut) on a per-particle basis and as efficiently as the original VSV-adapted GPC bearing all six substitutions [rVSV-CCHFV GPC(6-Mut)] (Fig 4B). These findings indicate that L518V and L1638R are minimally sufficient for rVSV-CCHFV GPC rescue and infection, and they reinforce the conclusion that D525G and Y728S contribute to rVSV fitness whereas S386T and R1648Q do not.

**Figure 4.**
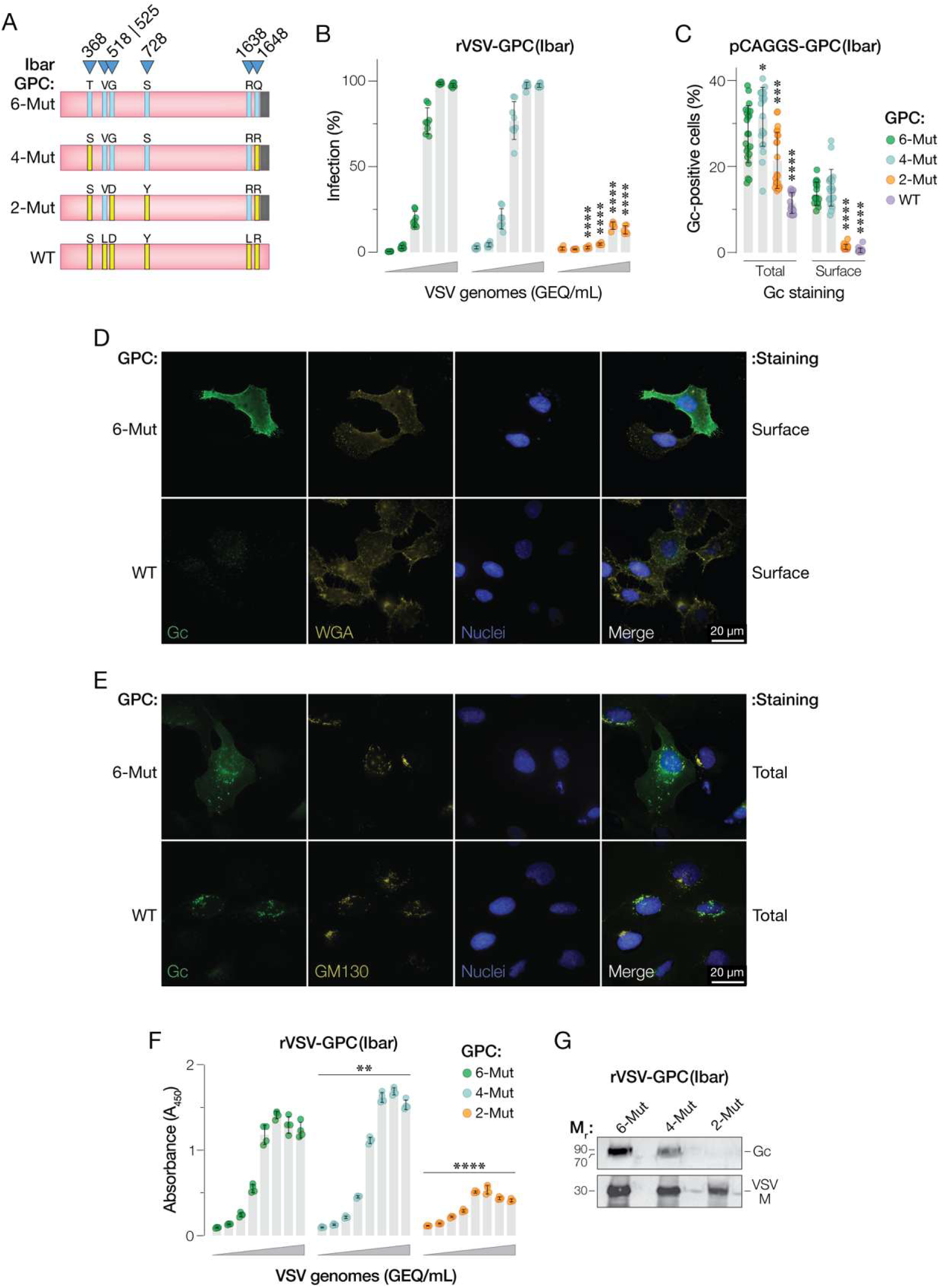
Identification and characterization of GPC substitutions optimally and minimally required for VSV adaptation. **(A)** GPC constructs carrying all six amino acid substitutions identified in rVSV-CCHFV GPC (6-Mut), the four essential substitutions (4-Mut) or the two minimal substitutions (2-Mut) are shown. Yellow and blue rectangles indicate WT and mutant residues, respectively, at the indicated positions. **(B)** rVSVs bearing the GPCs shown in **(A)** were rescued and genome-normalized preparations were exposed to Vero cells. GEQ, genome equivalents. Infection was scored by mNg expression at 16-18 h post-infection. Means ± s.d. from three independent experiments (*n*=8) are shown. Groups (6-Mut vs. others) were compared by two-way ANOVA with Tukey’s post hoc test. (C) GPCs shown in (A) were expressed from plasmids in U2OS cells, and the percentage of cells positive for Gc in each imaged field (≈X cells) was enumerated by immunofluorescence microscopy of permeabilized (Total) and intact cells (Surface). Means ± s.d. from four independent experiments (n=22) are shown. Groups (6-Mut vs. others) were compared by two-way ANOVA with Dunnett’s post hoc test. (D–E) Subcellular localization of rVSV-adapted GPC. U2OS cells were transfected with GPC, and intact and permeabilized cells were visualized in (D) and (E), respectively. GPC was stained with Gc-specific mAb ADI-36145. Wheat germ agglutinin (WGA) and GM130 were used as markers of the plasma membrane and cis-Golgi network, respectively. (F) The indicated genome-normalized rVSV-CCHFV GPC preparations were coated onto ELISA plates, and GPC was quantified with Gc-specific mAb, ADI-36145. A450, absorbance at 450 nm. Means ± s.d. from two independent experiments (n=4) are shown. The area under the curve (AUC) was calculated for each set of serial dilutions was calculated, and groups (6-Mut vs. others) were compared by one-way ANOVA with Dunnett’s post hoc test. G. Western blot analysis of genome equivalents of the indicated rVSVs for CCHFV Gc and the VSV matrix protein (M) with mAbs: ADI-36145 and Polyclonal VSV-M targeting Rabbit supp. respectively. M_r_, relative molecular weight. ** p < 0.01; *** p < 0.001; **** p < 0.0001. Only the statistically significant comparisons are shown.

To begin to investigate the mechanism(s) by which these suites of substitutions drive rVSV-CCHFV GPC infection and multiplication, we transiently transfected U2OS cells with plasmid constructs expressing GPC(2-Mut) and GPC(4-Mut) and compared their expression levels in permeabilized cells relative to WT and GPC(6-Mut) controls by immunofluorescence microscopy with a Gc ectodomain-specific mAb, ADI-36145 (Fig 4C-E). All the mutant constructs expressed Gc at higher levels than GPC(WT). More striking, the immunostaining of non-permeabilized cells with a Gc ectodomain-specific monoclonal antibody revealed that a substantial fraction of total Gc in GPC(4-Mut)- and GPC(6-Mut)-transfected cells, but not those of GPC(2-Mut) and GPC(WT), localized to the cell surface (Fig 4C-D). Consistent with these findings, Gc in GPC(WT) localized largely to the cis-Golgi network as expected, with no evidence of the diffuse staining observed with GPC(6-Mut) (Fig 4E). This behavior was mirrored by the incorporation levels of these GPC proteins into genome-normalized rVSV particle preparations [GPC(4-Mut) ≈ GPC(6-Mut) > GPC(2-Mut)], as determined by VSV Gc-specific ELISA (Fig 4F) and western blot (Fig 4G). Together, these results strongly suggest that D525G and/or Y728S contribute to rVSV-CCHFV GPC fitness at least in part by boosting the viral incorporation of GPC. Further, and concordant with the identification of ER retrieval motifs in the Gn and Gc C–tails by Gautam and co-workers, our findings point to roles for Y728S (Gn C–tail), L1638R (Gc C–tail), and the C–tail Δ14 deletion in relocalizing the VSV-adapted GPC to the cell surface as a likely mechanism for enhanced GPC incorporation into VSV particles [31].

### Cognate amino acid substitutions in a divergent CCHFV GPC afford successful VSV rescue

To assess if these infectivity- and incorporation-enhancing substitutions in IbAr GPC (Clade III) were transferable to an alternative CCHFV strain with a divergent viral genetic background, we attempted to generate replication-competent mCherry-reporter-expressing VSV bearing suites of these substitutions for a divergent GPC gene (Kosova-Hoti [K-Hoti], Clade V) [32] (Fig 5 and S1 Fig). rVSVs bearing K-Hoti GPC(4-Mut) and GPC(2-Mut) were successfully recovered and characterized as above. Similar to its IbAr GPC counterpart, K-Hoti GPC(4-Mut) afforded enhanced rVSV-CCHFV GPC infectivity relative to GPC(2-Mut) (Fig 5B). Further, we observed increases in the total and cell-surface expression of K-Hoti Gc in GPC for both mutants relative to WT in transfected cells, with 4-Mut conferring enhanced GPC localization at the cell surface relative to 2-Mut (Fig 5C-D). As seen for IbAr GPC(WT), K-Hoti GPC(WT) predominantly localized to the cis-Golgi (Fig 5E). Concordantly, and as with IbAr GPC, 4-Mut afforded enhanced K-Hoti GPC incorporation into VSV particles relative to 2-Mut (Fig 5F-G), reinforcing the contributions of the D525G and Y728S substitutions to this phenotype. We conclude that the substitutions identified in IbAr GPC reflect adaptations for altered expression, subcellular localization, and/or function that impact conserved structural and mechanistic features of the CCHFV entry glycoproteins.

**Figure 5.**
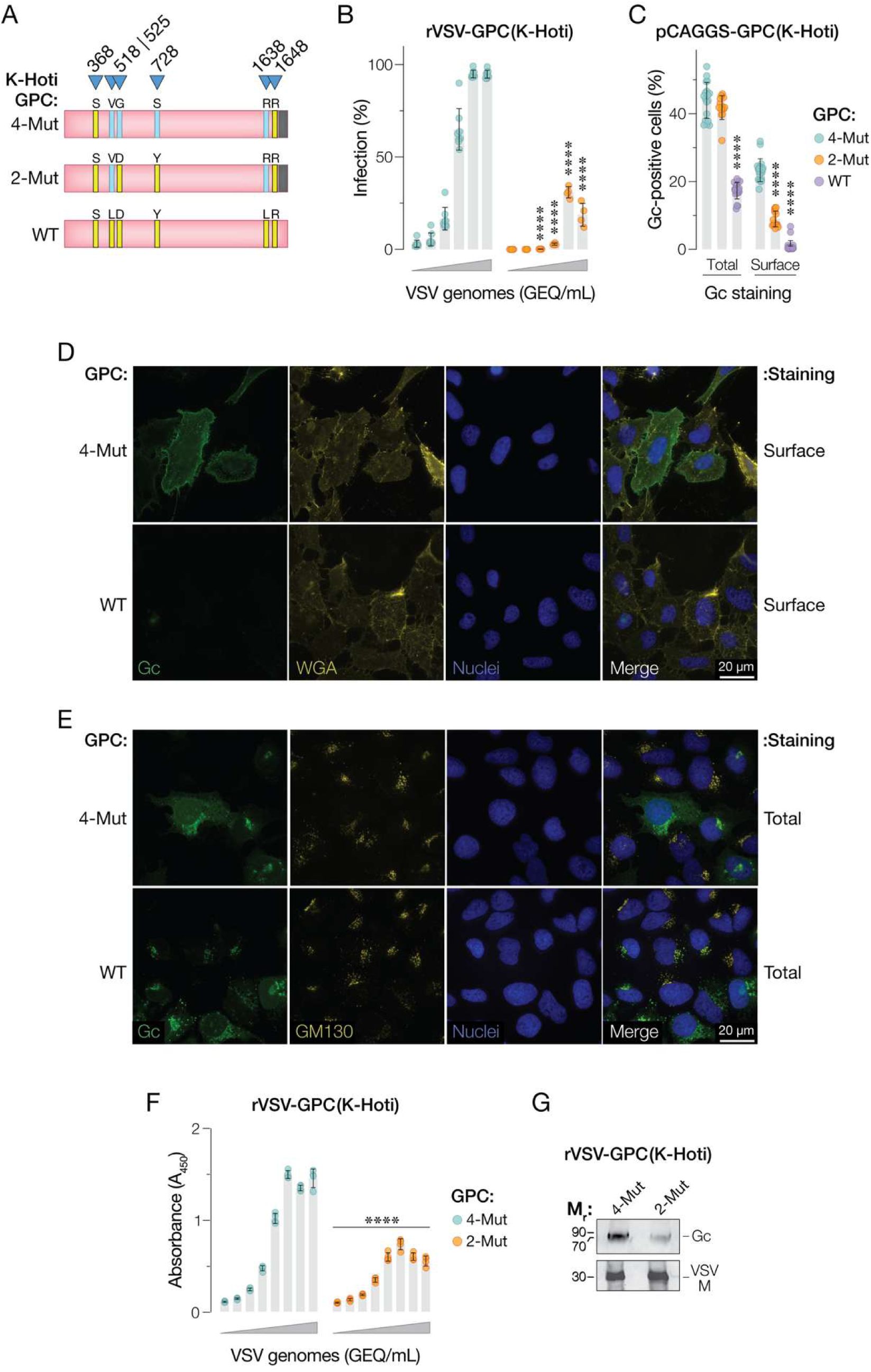
Identification and characterization of GPC substitutions required for rescue and replication of an rVSV bearing K-Hoti GPC. **(A)** K-Hoti GPC constructs carrying all four essential amino acid (4-Mut) or the minimal set of two substitutions (2-Mut) substitutions identified in rVSV-CCHFV GPC(IbAr) are shown. Yellow and blue rectangles indicate WT and mutant residues, respectively, at the indicated positions. **(B)** rVSVs bearing the GPCs shown in **(A)** were rescued and genome-normalized preparations were exposed to Vero cells. Infection was scored by mCherry expression at 16-18 h post-infection. Means ± s.d. from three independent experiments (n=8) are shown. Groups (4-Mut vs. others) were compared by two-way ANOVA with Tukey’s post hoc test. (C) GPCs shown in (A) were expressed from plasmids in U2OS cells, and the percentage of cells positive for Gc in each imaged field (≈X cells) was enumerated by immunofluorescence microscopy of permeabilized (Total) and intact cells (Surface). Means ± s.d. from four independent experiments (n=22) are shown. Groups (4-Mut vs. others) were compared by two-way ANOVA with Dunnett’s post hoc test. (D–E) Subcellular localization of rVSV-adapted GPC(K-Hoti). U2OS cells were transfected with GPC, and intact and permeabilized cells were visualized in (D) and (E), respectively. GPC was stained with Gc-specific mAb ADI-36145. Wheat germ agglutinin (WGA) and GM130 were used as markers of the plasma membrane and cis-Golgi network, respectively. (F) The indicated genome-normalized rVSV-CCHFV GPC preparations were coated onto ELISA plates, and GPC was quantified with Gc-specific mAb, ADI-36145. A450, absorbance at 450 nm. Means ± s.d. from two independent experiments (n=4) are shown. The area under the curve (AUC) was calculated for each set of serial dilutions and groups (4-Mut vs. others) were compared by one-way ANOVA with Dunnett’s post hoc test. G. Western blot analysis of genome equivalents of the indicated rVSVs for CCHFV Gc and the VSV matrix protein (M) with mAbs: ADI-36145 and Polyclonal VSV-M targeting Rabbit supp respectively. Mr, relative molecular weight. ** p < 0.01; *** p < 0.001; **** p < 0.0001. Only the statistically significant comparisons are shown.

### Substitutions at the GP38-Gn boundary influence GPC subcellular localization and VSV incorporation by modulating SKI-1 cleavage

Our findings pointed to a critical role in VSV adaptation for the L518V substitution in the SKI-1 cleavage site encoded in a sequence between GP38 and Gn. Recent work by other investigators also identified substitutions at or near this site during rVSV-CCHFV GPC adaptation [25, 33]. However, the mechanism(s) by which these substitutions enhance rVSV-CCHFV GPC infectivity remain poorly understood. To begin to selectively dissect the molecular basis of L518V’s effects, we generated single-cycle VSVs (scVSVs) bearing WT and VSV-adapted (6-mut) GPCs and a GPC in which only residue 518 was reverted from V to L in the VSV-adapted background to reconstitute the WT SKI-1 cleavage site (5-mut-RR<u>L</u>L) (Fig 6A). As expected, GPC(6-Mut) afforded a substantial increase in VSV per-particle infectivity relative to GPC(WT). Further, and in agreement with the forward-genetic studies in Fig 3, the V518L reversion in GPC(5-mut-RRLL) was sufficient to completely abrogate the infectivity advantage of GPC(6-Mut), despite the presence of the other adaptive substitutions (Fig 6B). This loss of viral infectivity was accompanied by a corresponding reduction in the incorporation of GPC into VSV particles, as determined by anti-Gc mAb ELISA (Fig 6C). Indeed, scVSV preparations bearing WT, 6-Mut, and 5-mut-RRLL GPCs displayed similar infectivity when normalized for Gc content instead of viral genome content (Fig 6D), suggesting that the L518V substitution primarily influences the steady state levels of pre-fusion GPC and/or its cell-surface localization, at least in the context of the other VSV-adaptive substitutions.

**Figure 6.**
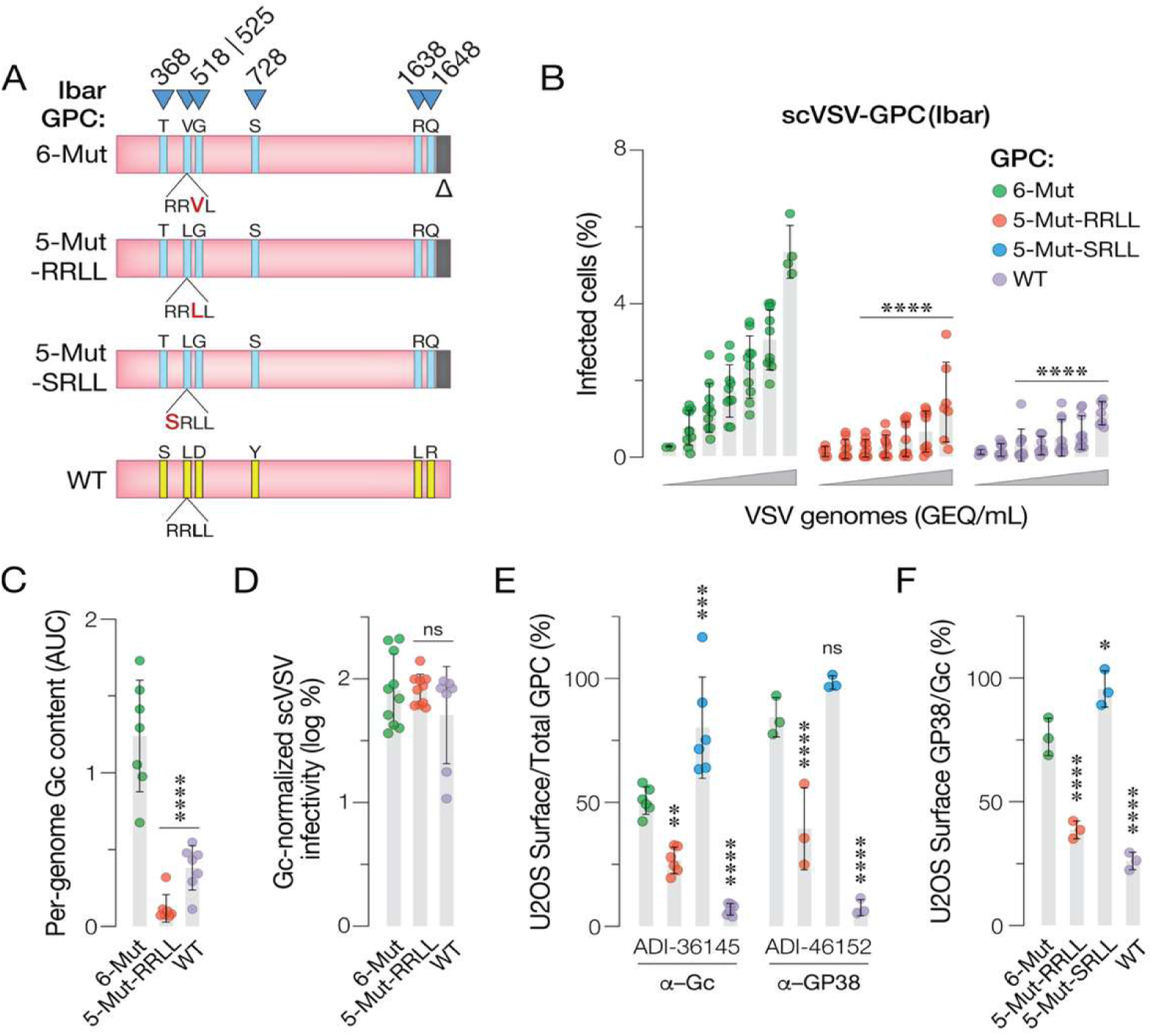
Effects of substitutions at the GP38-Gn cleavage site on VSV-GPC infectivity, GPC subcellular localization, and GPC processing. (A) GPC(IbAr) constructs carrying the indicated substitutions are shown. Yellow and blue rectangles indicate WT and mutant residues, respectively, at the indicated positions. (B) Single-cycle scVSVs bearing the GPCs shown in (A) were rescued and genome-normalized preparations were exposed to Vero cells. Infection was scored by mNg expression at 16-18 h post-infection. Means ± s.d. from three independent experiments (n=8) are shown. Groups (6-Mut vs. others) were compared by two-way ANOVA with Tukey’s post hoc test. (C) The indicated genome-normalized scVSV-GPC preparations were coated onto ELISA plates, and GPC was quantified with Gc-specific mAb, ADI-36145. A450, absorbance at 450 nm. Means ± s.d. from three independent experiments (n=10) are shown. The area under the curve (AUC) was calculated for each set of serial dilutions. Groups (6-Mut vs. others) were analyzed by one-way ANOVA with Dunnett’s post hoc test. (D) scVSVs normalized for Gc content in (C) were exposed to Vero cells, and infection was scored by eGFP expression as above. Means ± s.d. from three independent experiments (n=9) are shown. Groups ((6-Mut vs. others) were compared by two-way ANOVA with Tukey’s post hoc test. (E-F) GPCs were expressed from plasmids in U2OS cells, and the percentage of cells positive for Gc and GP38 were enumerated by immunofluorescence microscopy of permeabilized (Total) and intact cells (Surface) with the indicated mAbs. Ratios of the percentages of cells in each imaged field (≈X cells) with Surface vs. Total staining for GP38 and Gc (E) and Surface GP38 vs. Surface Gc staining (F) are shown. Means ± s.d. from three independent experiments (n=3–6) are shown. Groups ((6-Mut vs. others) were compared by two-way ANOVA with Šídák’s post hoc test. ns p > 0.05; ** p < 0.01; *** p < 0.001; **** p < 0.0001.

To directly examine the subcellular distribution of GPC at steady state, we next transfected constructs encoding these GPC variants into U2OS cells, stained the intact cells with Gc- and GP38- specific mAbs, imaged them by IF, and quantified Gc and GP38 staining at the cell surface (Fig 6E). When normalized to total Gc and GP38 staining in permeabilized cells, respectively, Gc and GP38 were expressed at much higher levels at the cell surface for GPC(6-Mut) than for WT, as expected (Fig 6E; also see Fig 4). Moreover, reversion of 518V to L in GPC(5-mut-RRLL) completely abrogated this enhancement in the cell-surface localization of Gc and GP38 (Fig 6E).

We next postulated that, if restoration of the SKI-1 site reduces GPC localization at the cell surface, then the complete loss of the SKI-1 site might have the opposite effect—a further increase in GPC relocalization. To test this hypothesis, we transfected U2OS cells with a GPC construct encoding GPC(5-mut-<u>S</u>RLL). As predicted, destruction of the SKI-1 consensus site afforded enhanced localization of Gc to the cell surface relative to GPC(6-mut) (Fig 6E). Further, 5-mutant GPC constructs encoding RRVL, SRLL, and RRLL displayed levels of relative GP38 staining at the cell surface that were concordant with the expected effects of these genotypes on GP38-Gn cleavage by SKI-1 (SRLL > RRVL > RRLL ≈ WT) (Fig 6F)[34]. We conclude that the substitution L518V enhances steady state levels of the VSV-adapted GPC at the cell surface by reducing (but not eliminating) GP38-Gn cleavage by SKI-1 in the secretory pathway.

### rVSV-optimized substitution at the SKI-1 site influences cell entry and modulates the pH threshold of membrane fusion by CCHFV virus-like particles

The preceding experiments were performed with rVSVs bearing CCHFV GPC. To examine the effects of the SKI-1 site substitutions in a setting more relevant to the biology of CCHFV, we generated transcription-and-entry-competent virus-like particles (tecVLPs) encoding a luciferase reporter gene and bearing WT and SKI-1 mutant GPCs, as described previously [35] (Fig 7A). Vero cells were then exposed to genome-normalized preparations of these tecVLPs, and luciferase reporter activity was used as a measure of entry efficiency. GPC(SRLL) displayed no tecVLP reporter activity relative to WT, pointing to the importance of SKI-1 cleavage at the GP38-Gn boundary for CCHFV entry. Unexpectedly, however, GPC(RRVL) afforded a ≈500% increase in tecVLP entry relative to WT (Fig 7B).

**Figure 7.**
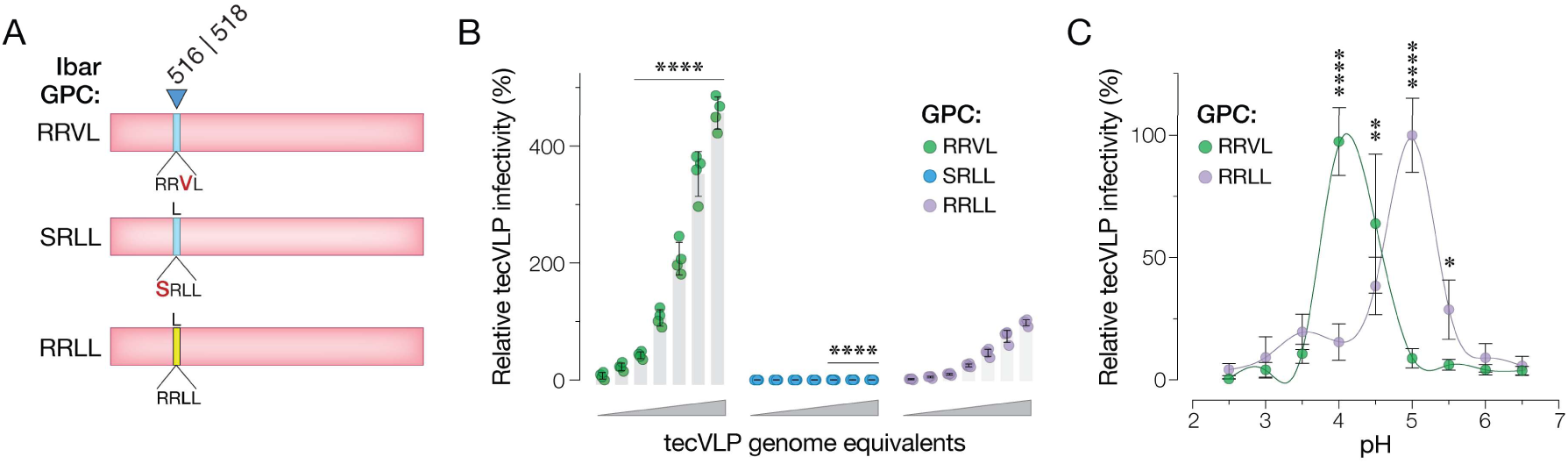
Generation and characterization of tecVLPs bearing substitutions at the GP38-Gn cleavage site. (A) GPC(IbAr) constructs carrying the indicated substitutions are shown. Yellow and blue rectangles indicate WT and mutant residues, respectively, at the indicated positions. (B) Minigenome-normalized tecVLPs generated from the GPC constructs in (A) were exposed to Vero cells. Infection was scored by luciferase activity at 16-18 h post-infection. Means ± s.d. from two independent experiments (n=3) are shown. Groups (RRLL vs. others) were compared by two-way ANOVA with Tukey’s post hoc test. (C) The activity of the indicated tecVLPs in a “fusion-from-without” infection assay in Vero cells. Means ± 95% confidence intervals from four independent experiments (n=18) are shown. Groups (RRLL vs. RRVL) were compared by two-way ANOVA with Šídák’s post hoc test. ** p < 0.01; *** p < 0.001; **** p < 0.0001. Only the statistically significant comparisons are shown.

Maturational or activating cleavage of glycoproteins (Class I) or their partner proteins (Class II) is often associated with structural rearrangements that are required for viral membrane fusion but that conversely render the glycoproteins more susceptible to premature triggering and inactivation[15, 36–39]. We postulated that this is also the case for GP38-Gn cleavage by SKI-1, and that partial inhibition of cleavage in GPC(RRVL) might stabilize its pre-fusion conformation, thereby enhancing tecVLP (and rVSV) infectivity. We and others have shown that CCHFV GPC-mediated cell entry requires endosomal acidic pH [40, 41]. Given that changes in the pH threshold for membrane fusion are a hallmark of structural changes that alter viral glycoprotein stability, we compared the pH thresholds of tecVLPs bearing WT and GPC (RRVL) in a “fusion from without” assay [42]. WT tecVLPs displayed a pH optimum of ≈5.5 (Fig 8C), as we showed previously in Wang & Monticelli et al. (2025). Remarkably, RRVL lowered GPC’s optimum fusion pH by an entire pH unit: ≈70% of entry events occurred between pH 4.5–6.5 for GPC(WT) but only ≈20% of entry events for GPC(RRVL) (Fig 7C). These findings point to currently undescribed structural changes associated with SKI-1 cleavage at the GP38-Gn boundary that influence the stability of GPC pre-complexes and regulate fusion triggering. Further, they help rationalize the recurrent selection of GPC mutants with a defective (but partially functional) SKI-1 cleavage site during rVSV-CCHFV GPC adaptation.

**Figure 8.**
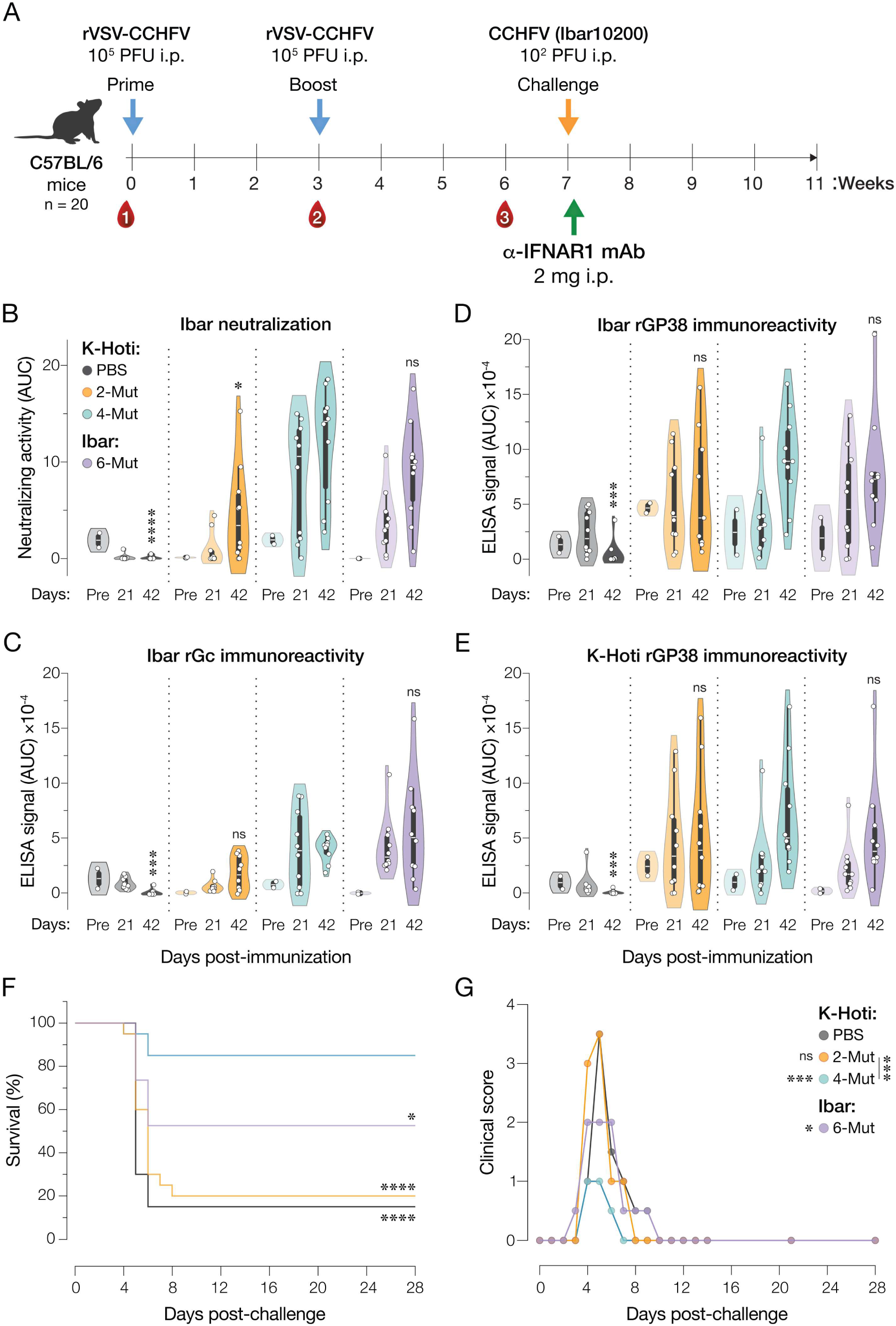
Protective efficacy of rVSVs bearing K-Hoti and IbAr GPC with distinct suites of adaptive substitutions. (A) Design of rVSV-CCHFV GPC immunization and CCHFV challenge study in a transiently immunosuppressed model of CCHFV challenge. (B) Capacity of murine immune sera from (A) to neutralize CCHFV(IbAr10200) infection. Violin plots are shown for area under the curve (AUC) calculated from each neutralization curve (n=2 mice for pre-immune controls [Pre], n=10 mice for immunized groups [days 21 and 42 post-immunization)]. White line, median. Thick center bar, interquartile range. (C–E) Immunoreactivity of the murine sera from (A) against the indicated recombinant GPC-derived proteins was determined by ELISA. Data were analyzed as in (B). In (B–E), groups (day 42; K-Hoti 4-Mut vs. others) were compared by two-way ANOVA with Tukey’s post hoc test. (F) Survival following CCHFV challenge (n=10 mice per group) was analyzed by two-sided Mantel-Cox (log-rank) tests. (G) Clinical scores of animals in (F) were compared (PBS vs. others, unless otherwise shown) by two-way ANOVA with Tukey’s post hoc test. ns p > 0.05; ** p < 0.01; *** p < 0.001; **** p < 0.0001.

### rVSV bearing K-Hoti GPC(4-mut) confers enhanced protection over IbAr GPC(6-mut) in a lethal murine model of heterologous CCHFV challenge

Previous work has shown that rVSVs bearing GPC from the prototypic CCHFV IbAr strain with distinct suites of adaptive substitutions can afford varying degrees of antiviral protection in lethal murine challenge models [19]. However, the vaccine efficacy of GPCs from divergent viral strains remains unexplored. Accordingly, we assessed the capacity of rVSVs bearing IbAr GPC(6-Mut), the minimally adapted K-Hoti GPC(2-Mut), or the fully VSV-adapted K-Hoti GPC(4-Mut) (see Fig 5) in their ability to protect against lethal CCHFV challenge in a transiently immunosuppressed murine model. First, C57/BL6 mice were subjected to prime-and-boost immunizations with equivalent infectious doses (1×10^5^ PFU) of each rVSV-CCHFV GPC (Fig 8A). Serological analysis at days -1 (pre-immune), 21 (post-prime), and 42 (post-boost) provided evidence for a robust neutralizing antibody response in the fully adapted VSV-GPC groups and a measurable but reduced response in the minimally adapted K-Hoti GPC(2-Mut) group (Fig 8B). A trend towards higher serological reactivity, most notably to the heterologous (IbAr) Gc and GP38 antigens, was also observed in the fully adapted rVSV-GPC groups but did not reach statistical significance (Fig 8C-E).

Immunized mice were challenged with a lethal dose of CCHFV(IbAr) and then administered α-IFNAR1 mAb (Clone MAR1-5A3) 24 h post challenge to effect immunosuppression, as described previously[22]. Animals were monitored for morbidity and mortality for 28 days (Fig 8F-G). Most of the animals succumbed to viral challenge in the PBS vehicle group. Little or no protection was observed in the K-Hoti GPC(2-Mut) group. By contrast, IbAr GPC(6-Mut) afforded partial protection at this vaccine dose, concordant with previous findings with rVSVs bearing partially and fully adapted IbAr GPC. Most strikingly, the fully adapted K-Hoti GPC(4-Mut) afforded near-complete protection (Fig 8F). Trends in clinical scoring (Fig 8G) mirrored the protection results. Thus, despite the administration of equivalent rVSV inoculum, the fully adapted rVSV-CCHFV GPC vectors were more protective against heterologous viral challenge than their partially adapted counterparts.

## Discussion

In this study, we investigated the molecular basis of adaptation of CCHFV GPC to support high levels of viral replication by an rVSV-CCHFV GPC vaccine vector previously described by Rodriguez *et al.* [19]. We identified a suite of four amino acid substitutions in GPC that is necessary and sufficient for rVSV-CCHFV GPC adaptation. These substitutions could be transferred to a divergent CCHFV GPC, and an rVSV-CCHFV GPC bearing these substitutions could protect against heterologous virus exposure. Our mechanistic studies point to two distinct roles for these essential substitutions: two amino acid changes in the cytoplasmic tails of Gn and Gc relocalize the pre-fusion glycoprotein from the Golgi complex to the cell surface to enhance its incorporation into budding VSV virions. The remaining two amino acid changes at the GP38-Gn boundary in the GPC ectodomain appear to enhance expression of the pre-fusion glycoprotein complex and protect it from premature inactivation by SKI-1/S1P during intracellular trafficking. Our findings suggest a new model for GP38’s roles in the biogenesis, trafficking, and deployment of CCHFV GPC.

Previous work identified L1638R in the Gc C–tail as a GPC substitution associated with its adaptation to VSV [19]. Herein, we show that L1638R is critical for efficient rVSV-CCHFV GPC infection and multiplication (Fig 3D). We also uncovered a second, hitherto undescribed, substitution in the Gn C–tail, Y728S (Fig 3D), which confers a more subtle fitness benefit. Strikingly, Gautam and coworkers recently described and characterized tyrosine-based endocytic motifs (Yxxφ) at precisely these positions in the Gn and Gc C–tails (<u>Y</u>REL at residues 728–731 and YRH<u>L</u> at residues 1635–1638, respectively) [31]. Recognition of YXXLφ motifs (where φ is I,L,F,M,V) in the C–tails of transmembrane proteins by AP-2 adapter complexes mediates their sorting into clathrin-coated pits and subsequent internalization [43, 44]. Mutation of the Gc YRHL motif by these authors increased GPC localization at the cell surface; however, they did not observe a measurable effect upon mutation of Gn YREL [31]. The significant, if more nuanced, role for Y728S in rVSV-CCHFV GPC fitness suggests that Gn YREL does indeed contribute to GPC re-internalization from the plasma membrane, likely in concert with Gc YRHL. We conclude that loss of the Yxxφ endocytic motifs in the Gn and Gc C–tails, together with the 14-amino acid deletion at the Gc C–tail, which eliminates a non-canonical YXHXX recognition motif for COPI coatomer binding and ER retention [31, 45], serves to redistribute a substantial fraction of the GPC pre-fusion complexes to the plasma membrane at steady state, where they are available for incorporation into budding VSV virions. Our findings are consistent with previous work using single-cycle VSV pseudotypes and rVSVs, which showed that a larger deletion at the Gc C–terminus encompassing both the YHRL and YxHxx motifs enhances VSV-GPC infectivity [25, 46]. Interestingly, rVSV-CCHFV GPC does not contain substitutions in the acidic cluster and di-leucine sorting motifs in the Gn and Gc C–tails, which appear to be important for CCHFV VLP assembly and budding [31, 45]. Therefore, these motifs either do not interfere with rVSV-CCHFV GPC infectivity or are required for the biogenesis of pre-fusion glycoprotein complexes in a manner that remains to be defined.

Our forward-genetic studies revealed that two amino acid substitutions at the GP38-Gn boundary in the GPC ectodomain enhance rVSV-CCHFV GPC fitness. Both are located in a flexible or variably placed loop not visualized in the cryo-EM structure of a soluble, heteromeric GP38-Gn-Gc complex resembling the pre-fusion conformation of the CCHFV glycoprotein [16]. L518V modifies the consensus site for GPC cleavage by the proprotein convertase SKI-1, a key step in the proteolytic maturation of GPC during glycoprotein assembly (further discussed below) [14, 47]. D525G was uncovered in this study and has not been described previously. This substitution’s structural and functional effects are unclear at present, but it may serve to modulate GP38-Gn cleavage by SKI-1, alter the quaternary packing of GP38-Gn-Gc structural units on the virion surface, or both.

We found that the L518V substitution at the GP38-Gn boundary, which modifies the consensus SKI-1 cleavage site (RR<u>L</u>L→RR<u>V</u>L) is indispensable for rVSV-CCHFV GPC infectivity. Mutations at the SKI-1 cleavage site are a recurrent theme in previous work with VSVs bearing CCHFV GPC; however, the molecular bases of their effects on VSV infectivity are unknown [19, 25]. Here, we show that <u>either</u> restoring the consensus SKI-1 cleavage site found in WT (RR<u>V</u>L→RR<u>L</u>L) or disrupting it completely (<u>R</u>RLL→<u>S</u>RLL) in the background of the other VSV-adaptive substitutions is highly deleterious to VSV infectivity. Previous work on the specificity of SKI-1 cleavage indicates that RRLL→RRVL→SRLL should progressively reduce GP38-Gn processing [34, 48], a prediction borne out by the ratio of GP38 to Gc measured at the cell surface with each GPC (RRLL > RRVL > SRLL). We infer that L518V is a “Goldilocks” substitution that enhances VSV infectivity by balancing the opposing effects of GP38-Gn cleavage on GPC cell-surface localization (<u>S</u>RLL >> <u>R</u>RLL) and GPC-mediated cell entry (<u>R</u>RLL >> <u>S</u>RLL).

To uncover the specific contribution of the SKI-1 site substitutions to GPC cell-surface localization and VSV infectivity vis-à-vis the other adaptive changes, we investigated the behavior of GPCs in which only the SKI-1 site was modified. Unsurprisingly, replacing the WT SKI-1 site (RRLL) with RRVL or SRLL did not re-localize GPC to the cell surface in the absence of the C–tail mutations in Gn and Gc. Thus, modulating GP38-Gn cleavage alters the structure of the GPC ectodomain in a manner necessary but not sufficient for GPC relocalization. To determine if these structural changes independently alter GPC’s functional behavior, we tested the capacity of GPC(RRVL) and GPC(SRLL) to generate CCHFV tecVLPs. As expected, tecVLPs bearing GPC(SRLL) were poorly infectious. In striking contrast, tecVLPs bearing GPC(RRVL) were even more infectious than their WT counterparts. Given that maturational or activating cleavages in the partner proteins of Class II fusion glycoproteins are often associated with structural rearrangements required for viral membrane fusion, we postulated that inhibition of some GP38-Gn cleavage events in particles bearing multimeric GPC(RRVL) complexes might alter their fusogenicity[36, 37, 49]. In agreement with this hypothesis, we measured a downward shift of ≈1 pH unit in the acidic fusion threshold of tecVLPs bearing GPC(RRVL) relative to those bearing GPC(WT). Indeed, inhibiting the furin cleavage of its p62 partner protein lowers the acidic fusion threshold of the Class II alphavirus glycoprotein (E1) to a similar extent [49–52]. Although more work is needed to uncover the precise molecular mechanism for this effect, our findings suggest a model in which GP38-Gn cleavage induces, or sets the stage for, structural changes that are required for the acid-triggered dissociation of Gn and Gc and fusion-related conformational Gc rearrangement during entry (S2 Fig). The timing of and trigger for GP38 dissociation from cleaved pre-fusion GPC are currently unknown.

Why might reducing the fusogenicity of GPC benefit both VSVs and tecVLPs? Freitas *et al.* provided evidence that GP38 is crucial for assembly of pre-fusion GPC complexes, and McFadden *et al.* demonstrated that GP38 is an integral component of this complex [11, 16]. Our work extends these findings by pointing to a key role for GP38 (and GP38-Gn cleavage) in protecting pre-fusion GPC complexes from premature triggering and inactivation in the acidic environment of the TGN (pH ≈6.0) and secretory vesicles (pH ≈5.5) [53] (S2 Fig). Such a role for GP38 would be directly analogous to that of E3 (and p62→E1+E2 cleavage) for alphavirus glycoproteins and pr (and prM→pr+M cleavage) for flavivirus glycoproteins [37, 54, 55]. We speculate that tecVLPs’ reduced sensitivity relative to VSVs in this regard (i.e., GPC(RRLL) can produce infectious tecVLPs but not VSVs) ensues from their distinct intracellular sites of virion assembly. Specifically, tecVLPs, like authentic CCHFV, bud into the Golgi lumen, whereas VSVs bud at the plasma membrane [35, 56]. It is conceivable, therefore, that multimeric GPC complexes incorporated into CCHFV virions in the Golgi are more resistant to acid-induced inactivation upon cleavage than the isolated GPC complexes that must traffic to the cell surface with their pre-fusion conformation intact prior to incorporation into budding VSVs. The premature inactivation of trafficking GPC complexes in the secretory pathway would also explain the greatly reduced amounts of pre-fusion GPC localized to the cell surface in VSV-adapted GPC(RRLL), as is observed with alphaviruses: mutations in the E3 protector protein that disrupt its interaction with the fusion glycoprotein E1 prevent delivery of the pre-triggered E1 molecules to the cell surface [37].

Finally, we found that the four key amino acid substitutions identified in this study were necessary and sufficient to generate a highly replication-competent rVSV bearing GPC from the Clade V CCHFV strain, K-Hoti. Although the two essential substitutions we mapped with IbAr GPC (L1638Q and L518V; IbAr numbering) were minimally sufficient for rVSV rescue, inclusion of the other substitutions (D525G and Y728S) maximized GPC cell-surface localization and virion incorporation. Accordingly, the fully optimized rVSV-CCHFV GPC(K-Hoti-4-Mut) afforded the highest levels of protection against heterologous CCHFV(IbAr) challenge in a transiently immunosuppressed murine model, even improving upon the optimized rVSV-CCHFV GPC(IbAr-6-Mut). The superior performance of rVSV-CCHFV GPC(K-Hoti-4-Mut) as a vaccine vector relative to its minimally adapted counterpart, rVSV-CCHFV GPC(K-Hoti-2-Mut), may stem from enhancements in one or more of the following: the levels of (i) cell-surface GPC expressed in infected cells; (ii) budded virus-like particles released from infected cells; (iii) GPC in budded VSV particles; and (iv) infected cell and/or vector loads due to improved VSV replicative fitness; and (v) inherent differences in the immunogenicity of the GPC molecules. The molecular basis of enhanced protection is thus likely to dictate if these GPC substitutions can improve the performance of GPC-based immunogens delivered via other vaccine modalities, such as RNA and proteins. These studies are currently underway, as are experiments to determine which, if any, of these CCHFV substitutions are transferable to other orthonairoviruses. In this regard, it is interesting to note that the GPC N–tail and C–tail sequences from many of the latter viruses appear to lack some or all of the tyrosine-based endocytic motifs found in CCHFV, suggesting they employ distinct mechanisms to regulate GPC intracellular trafficking. The identification and perturbation of these mechanisms may provide new approaches to generate and optimize “plug-and-play” VSV vaccine vectors targeting orthonairoviruses.

## Author Contributions

Conceptualization: J.B., S.R.M., A.S.W., A.S.H., K.C.

Investigation: J.B., S.R.M., P.S., C.A.C., T.G.B., A.W., E.M., R.R.B., M.M.S., A.S.W., D.W.H., K.C.

Methodology: J.B., S.R.M., P.S., A.W., E.M., M.M.S., A.S.W., R.W.C., D.W.H.

Supervision: T.W.G., J.H.E., J.M., A.S.H., K.C.

Visualization: J.B., S.R.M., P.S., E.M., K.C.

Writing – original draft: J.B., S.R.M., K.C.

## Acknowledgments

We would like to thank Estefania Valencia, Javier Janer, Karyme Paez, and Matt Ramirez for laboratory management; Ezgi Kasikci for valuable input; all the members of our laboratories; and members of the PROVIDENT consortium for technical advice and support. We would like to thank Kandis Cogliano and Cecilia O’Brien for laboratory management and programmatic support. We would also like to thank Christy Hjorth in the McLellan Laboratory for their help with the production of soluble CCHFV GPC. Lastly, the authors acknowledge the Flow Cytometry Core and its personnel at Albert Einstein College of Medicine and the following divisions at USAMRIID: the Veterinary Medicine Division and Statistics Division.

## Data Availability

All data associated with this study, including all raw data required to replicate the results, are present in the paper, its Supplementary files, and the files shared through public repositories (see below). The raw data generated in this study has been deposited in the Figshare database under accession code (10.6084/m9.figshare.c.8495061). Nanopore Sequencing data has been deposited in the Sequence Read Archive (SRA) under accession numbers (<u>PRJNA1271595</u>) and (<u>PRJNA1470180</u>) for initial sequencing and serial passage sequencing respectively.

## Disclaimers

The opinions, interpretations, conclusions, and recommendations presented are those of the authors and not necessarily endorsed by the U.S. Army or Department of Defense. The use of either trade of manufacturers’ names in this report does not constitute an official endorsement of any commercial products. This report may not be cited for purposes of advertisement. This does not constitute an endorsement by the U.S. government of this or any other contractors.

---

**Supplementary Fig 1.**
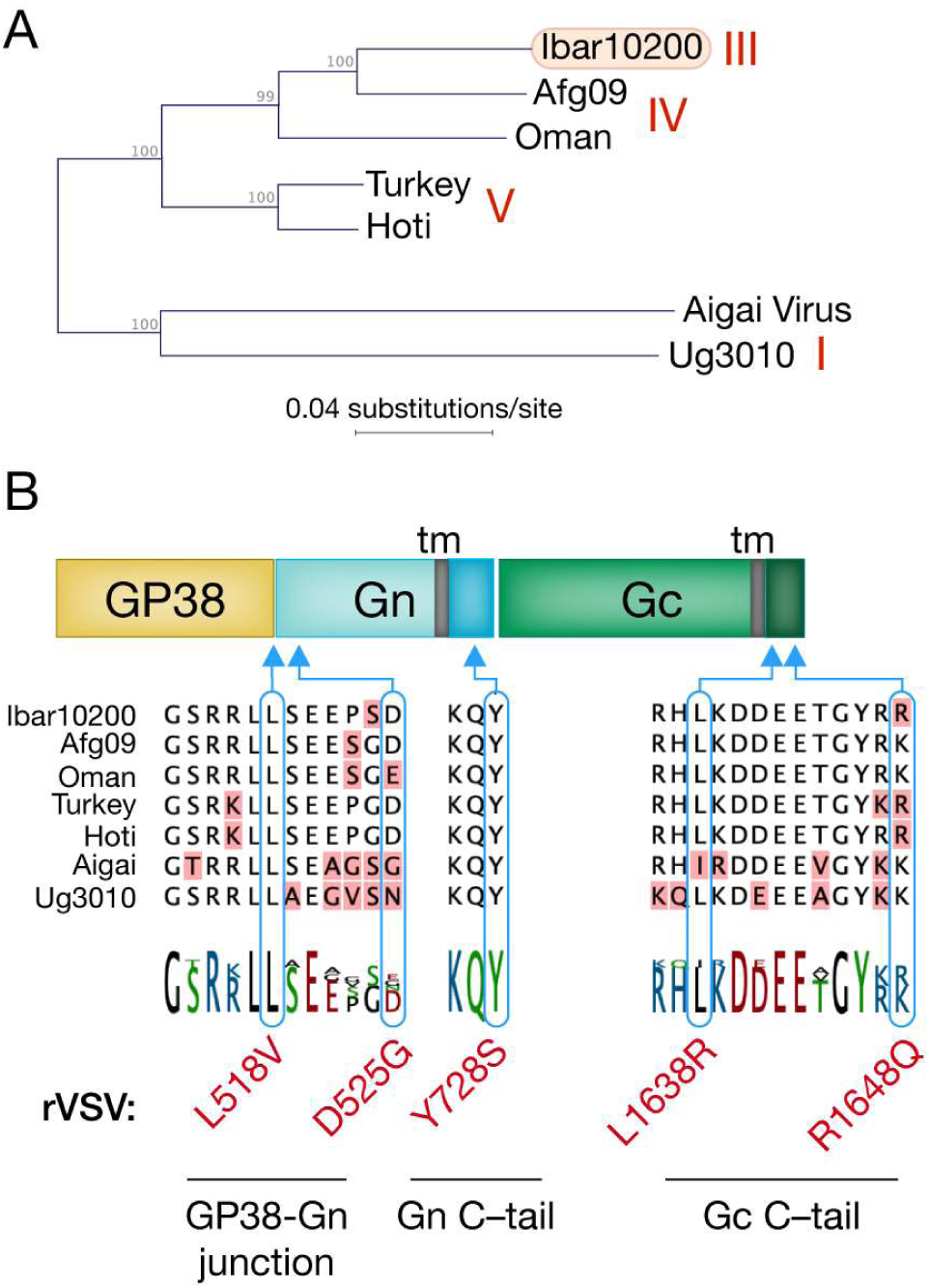
Conservation of GPC sequences at the positions mutated in rVSV-CCHFV GPC. A. Phylogenetic analysis of seven distinct CCHFV M segments across 4 relevant clades based on amino acid sequence. B. Global amino acid sequence alignment of the M segments from **(A)** highlighting the adaptive mutations found in rVSV-CCHFV GPC. Only the relevant GPC regions are shown.

**Supplementary Fig. 2.**
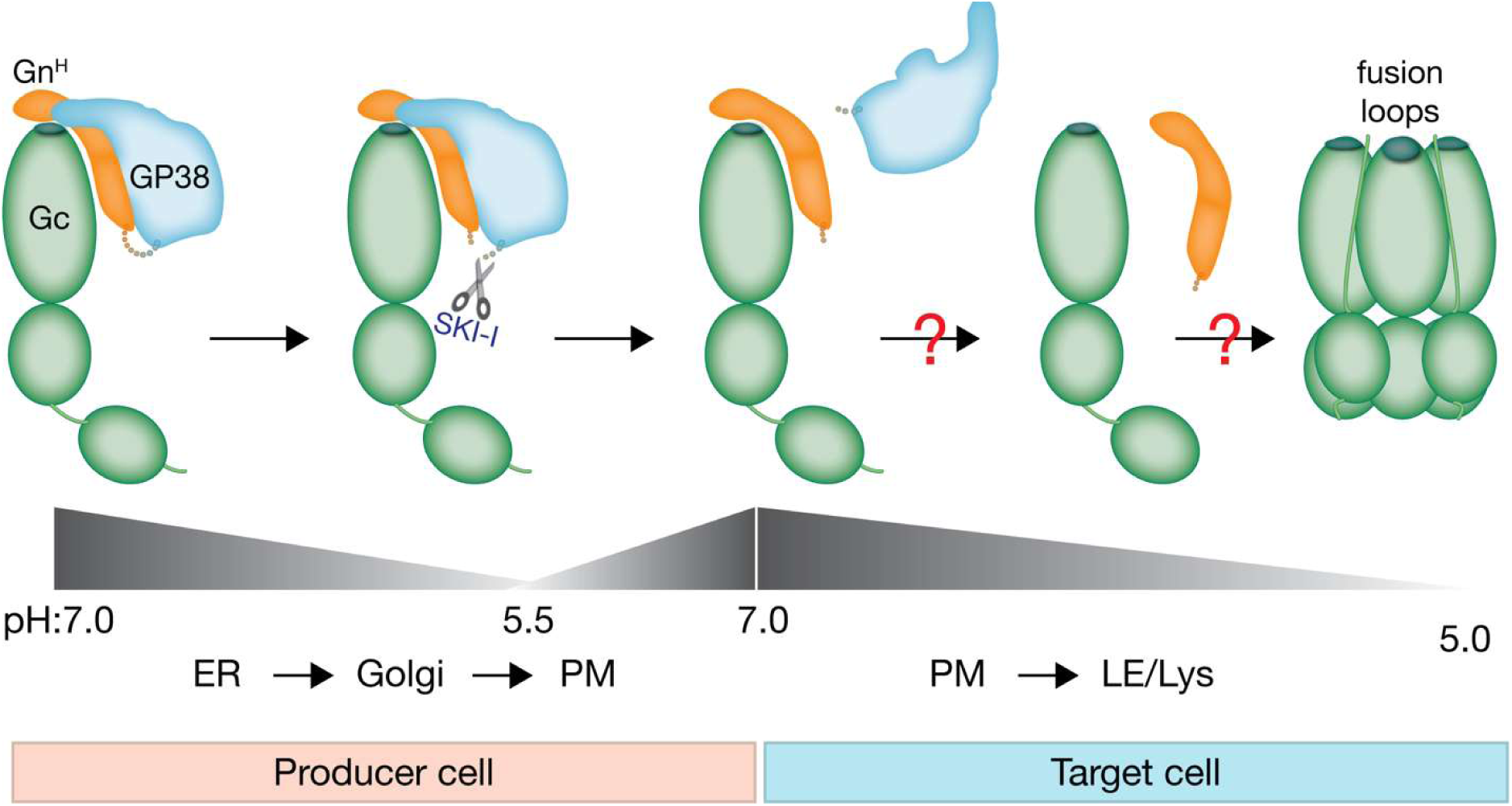
Speculative model of structural rearrangements in the GPC ectodomain during cellular egress and entry.

**Supplementary Fig. 3.**
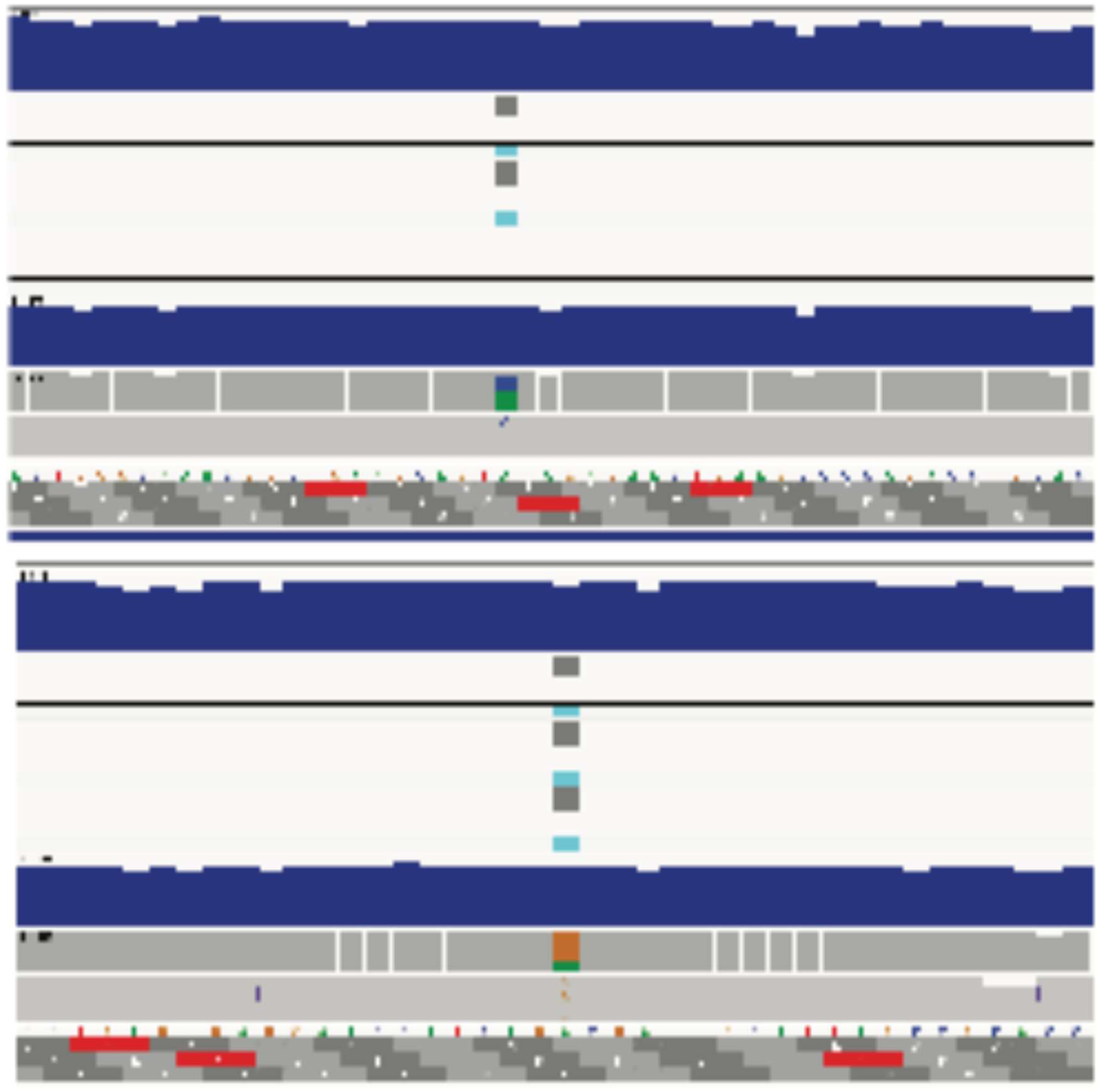
Analysis of GPC sequences from long-read sequencing of cDNA isolated from viral populations generated from rVSV rescue of rVSV-CCHFV GPC led to identification of D525G (A.) and Y728S (B.) mutations. See Materials and methods for analysis workflow, variant calls reported are in VCF format and visualized. Data from two pooled biological repeats are shown below.

## Materials and Methods

### Cell culture

African green monkey kidney Vero cells [American Type Culture Collection (ATCC), Cat# CCL-81] were cultured in high-glucose Dulbecco’s Modified Eagle Medium (DMEM; Gibco, Cat# 11965-092) supplemented with 2% heat-inactivated fetal bovine serum (FBS; Gemini Bio, Cat# 100-106, Lot A123000), 1% penicillin-streptomycin (Gibco, Cat# 15140-122), and 100 U/mL penicillin-streptomycin. Human osteosarcoma U-2 OS cells [ATCC, Cat# HTB-96] were maintained in McCoy’s 5A Medium Modified (Gibco, Cat# 16600082) supplemented with 10% FBS and 1% penicillin-streptomycin. Human hepatocarcinoma Huh 7.5.1 cells (a generous gift from C. M. Rice, The Rockefeller University, NY, and F. V. Chisari, The Scripps Research Institute, CA) were maintained in high-glucose DMEM (Life Technologies, Cat# 11965-092) supplemented with 10% FBS, 1% penicillin-streptomycin, and 1% non-essential amino acids (Life Technologies, Cat# 11140-050). Human embryonal kidney 293FT cells (ThermoFisher Scientific, Cat# R70007) were maintained in high-glucose DMEM supplemented with 10% FBS, 100 U/mL penicillin-streptomycin, and 25 mM HEPES (Life Technologies, Cat# 15630-080). A549 cells [ATCC, Cat# CCL-185] were maintained in DMEM/F-12 (Gibco, Cat# 11320-033) supplemented with 10% FBS (Gemini Bio, Cat# 100-106) and 1% penicillin-streptomycin.

### Recombinant VSV production and infection

To study CCHFV entry and infection in a BSL-2 setting, we utilized a replication-competent recombinant vesicular stomatitis virus (rVSV) expressing the glycoprotein complex (GPC) of CCHFV (rVSV-CCHFV GPC). This rVSV had previously been used for vaccine development and, for this reason, lacked a reporter [19]. To enable high-throughput screening and facilitate infection analysis, we generated a replication-competent reporter virus encoding a fluorescent mNeonGreen-tagged VSV phosphoprotein (P), termed rVSV-CCHFV GPC. To construct this virus, we used inverse PCR to clone the CCHFV GPC into a VSV backbone encoding mNg-P.

### Single cycle VSV production and infections

Single cycle VSV pseudotypes bearing CCHFV GPC and GFP reporters were generated by transfection as described previously [46]. 293FT cells were transfected with a pCAGGS expression vector containing the CCHFV GPC. Two days later cells were infected with a p2 stock of VSV(ΔG) pseudotyped with VSV G for 1 hour at 37 C. Cells were washed 8 times to remove residual VSV expressing G protein. Viral supernatants were harvested two days later and pelleted via ultracentrifugation at 20,000 xg in a SW-28 rotor for 2 h at 4 C. Viral preps were then analyzed via qPCR utilizing probes targeting VSV-M in the RNA genome, viral preps were also analyzed via ELISA to confirm presence of CCHFV GPC.

### VLP production and infections

Virus-like particles were produced as previously described [35]. Mutant GPCs were cloned into pCAGGS expression vectors. VLP titers were normalized via qPCR using primers targeting the minigenome expressed by VLP. (Probe: GATTACCAGTGTGCCATAGTGCAGGATCAC Primers: 5’-TAGTCGATCATGTTCGGCGT-3’ and 5’-ACCCTGTGGATGATCATCACT-3’). VLP infections were performed by infecting cells with genome copy normalized inputs of VLPs, cells were infected for 14 h at 37 C and 5% CO2. Cells were then lysed and developed using the Promega nanoluciferase system (Cat. N1120).

### Oxford nanopore sequencing

Oxford Nanopore long read sequencing was performed to identify mutations present in viral populations [57]. Viral RNA was isolated from supernatants harvested from infected cells. Supernatants were spun for 5 min at 5000 xg to pellet cells and subsequently viral RNA was isolated using the Zymo Quick-Viral RNA kit (Cat. R1035). Following RNA extraction viral RNA was reverse transcribed into DNA using the Invitrogen Superscript IV first strand synthesis system using primers:(CTCCAGCGGTATTGGCAGAT and CCACATCGAGGGAATCGGAA) (Cat. 18091050). Resulting cDNA was further amplified using Q5 HotStart Hi Fidelity polymerase with primers directly upstream and downstream of the CCHFV GPC (AGGCCTTAATGTTTGGCCTG and AAATCATTGAACTCGTCGGTCTC) (Cat. M0493S). Amplified cDNA was purified via gel extraction and prepped for long read analysis. Sample preparation was performed with Ampure XP beads (A63881) according to the protocol for direct sequencing of cDNA with barcoding (SQK-PCB114.24). Following purification of the library cDNA Nanopore libraries were sequenced with MinION MK1D device (MIN-101D) with Flow Cell (Cat. R10.4.1). Reads were aligned using minimap2, with a median depth of ∼2000x, structural variants were searched for using Sniffles, SNPs were called using NanoCaller, and genome assemblies were made using Flye [58–60].

### Immunofluorescence microscopy

Human U2OS osteosarcoma cells were plated on glass coverslips pre-coated with fibronectin. Cells were subsequently transfected with 500ng of pCAGGS expression vectors expressing CCHFV GPC. At 24 h post-transfection, cells were fixed for 5 min with 4% paraformaldehyde (ThermoFisher). Cells were then permeabilized with 0.1% Triton X-100 for 10 min at room temperature, then blocked with 10% FBS for 30 min. If cells were stained for surface expression levels of CCHFV GPC, they were not initially fixed or permeabilized and instead blocked on ice for 30 min with 10% cold FBS. After blocking, cells were incubated with either anti-Gc 36121 (1 µg/mL Adimab) or anti GP38-46152 (1 µg/mL Adimab) [27, 28]. In addition to CCHFV GPC targeting antibodies, anti GM130 Golgi markers (1:1000 Cell Signaling Cat. E9Z6S), anti-calreticulin ER markers (ThermoFisher Cat. MA5-15382) or Wheat Germ Agglutinin were used for cell surface staining (ThermoFisher Cat. W11261). Following staining for surface localization, cells were fixed with 4% paraformaldehyde. Secondary antibodies were then added, and slides were stained with Prolong mounting agent containing DAPI (ThermoFisher Cat. P36965) and imaged using a Zeiss Axio Observer inverted microscope with a 63x objective.

### ELISA

ELISA assays were performed by coating half area high binding plates (Corning Cat. CLS3694). VSV or VLPs were diluted in PBS prior to coating. Coated plates were then stored at 4 C overnight. Subsequently plates were washed three times with 0.1% PBS-Tween and blocked with 10% Milk for 1 hour at 37 C. Following blocking for 1 h, plates were washed three times with 0.1% PBS-Tween. Next, primary antibodies were added to the plate targeting either VSV-M (Polyclonal Sera Courtesy of Albert Einstein College of Medicine Hybridoma Core) for rVSV-CCHFV GPC or CCHFV GPC anti-Gc 36121 (1 µg/mL Adimab) for VLP ELISA, following addition of the antibody the plates were moved to 37 C for 1 hour[27, 28]. Next plates were further washed three times with 0.1% PBS-Tween. Plates were then coated with anti-human HRP secondary antibody (ThermoFisher A-10648) and incubated for 1 hour at 37 C. plates were washed three times with 0.1% PBS-Tween and developed using one step TMB peroxidase (ThermoFisher Cat. 34028). After developing for 5 min, Sulfuric acid was added and the plates were read on an Agilent Cytation 7.

### Western blotting

Virus like particles and VSV were analyzed by SDS-PAGE followed by western blotting for both GP38 and Gc. Cleavage patterns were analyzed by observing differences between wild type and mutant SKI-1 cleavage motifs. anti-Gc 36145 (1 µg/mL Adimab) and anti GP38-46152 (1 µg/mL Adimab) was used to detect CCHFV GPC in the VLP system [27, 28]. Input concentrations of VLPs were determined based on Minigenome targeting qPCR. Following primary antibody staining anti human HRP secondary antibody (ThermoFisher A-10648) was used to detect CCHFV GPC. For VSV based western blotting experiments VSV inputs were normalized based on qPCR targeting VSV-M. Anti-Gc 36145 (1 µg/mL Adimab) was used to detect GPC present on the surface of the VSV, anti-VSV-M polyclonal mouse sera were used to stain the samples. Secondary antibodies were then used: anti human HRP secondary antibody (ThermoFisher A-10648) and Goat anti-Mouse IgG (H+L) Cross-Adsorbed Secondary Antibody, Alexa Fluor™ 680 (ThermoFisher Cat. A-21057) were used. Blots were read on an Invitrogen iBright imager.

### Authentic virus enzyme-linked immunosorbent assay (ELISA)

ELISAs for each serum sample were run in duplicate. Flat bottom, high binding, half-area 96-well plates (Corning) were coated with 125 ng CCHFV rGc, sheep Fc-Tag (Native Antigen) or 75 ng recombinant GP38 protein[28] diluted in PBS at 4 °C overnight (∼18 h). The initial coating was decanted, and the plate was then blocked with 5% nonfat dry milk (BD Biosciences) diluted in PBS containing 0.05% Tween-20 (PBST) for 2 h at 37°C. Serial 3-fold dilutions were made for each mouse serum sample starting at a 1:20 dilution in blocking buffer and plates were loaded with dilutions in duplicate. Plates were incubated at ambient temperature for 2 h, washed three times with PBST, and incubated with horseradish peroxidase (HRP)-conjugated goat anti-mouse (1:2000 dilution; Jackson ImmunoResearch) diluted in blocking buffer for 1 hour at ambient temperature. Following incubation, plates were washed three times with PBST and developed at room temperature with 1-Step TMB Ultra Substrate (ThermoFisher). Reaction was stopped using 0.16 M sulfuric acid and absorbance was read at 450 nm wavelength, detected using a SpectraMax (Molecular Biosciences) microplate reader. Naïve sera collected prior to vaccination was used as an internal control for each group. A cutoff value was determined based on the average absorbance of the naïve control starting dilution plus 3 standard deviations. Only sample dilutions whose average were above this cut-off were registered as a positive signal. AUC values were computed for each ELISA curve (Baseline Y=0, Ignore peaks <10% of the distance from min to max Y). AUCs were log-transformed and compared for each serum and immunogen group by 2-way ANOVA with Tukey corrections for multiple comparisons. Statistical tests were performed in GraphPad Prism 10.0.3.

### Ethics statement for IbAr10200 *in vivo* challenge study

Murine challenge studies were conducted under Institutional Animal Care and Use Committee (IACUC)-approved protocols in compliance with the Animal Welfare Act, PHS Policy, and other applicable federal statutes and regulations. The facility where these studies were conducted (USAMRIID) are accredited by the Association for Assessment and Accreditation of Laboratory Animal Care, International (AAALAC) and adhere to the principles stated in the Guide for the Care and Use of Laboratory Animals, National Research Council, 2013. Humane endpoints were utilized during these studies and mice that were moribund, according to an endpoint score sheet and in line with IACUC-approved criteria, were humanely euthanized.

### IbAr10200 *in vivo* challenge study

4-week-old male and female C57BL/6J mice (strain #000664; The Jackson Laboratory) were vaccinated two times at 3-week intervals with 1x10^5^ plaque-forming units (PFU) of VSV or an equal volume of PBS vehicle control. Constructs were diluted in endotoxin-free PBS (ThermoFisher Scientific) to a total volume of 200 µL. Whole blood was collected prior to vaccination on days 0, 21, and 42 by submandibular bleed and sera were isolated from whole blood as described above. Mice were challenged on day 49. For the challenge, all mice were challenged with 100 PFU of CCHFV-IbAr10200 by the IP route. At 24 h post-challenge, mice were transiently immunosuppressed by treatment with 1.5 mg/mouse of mAb-5A3 (Leinco Technologies Inc.) via the IP route. Mice were monitored daily for weight changes, clinical score, and survival. Mice were scored on a 4-point grading scale; 1 defined by decreased grooming and ruffled fur, 2 defined by subdued behavior when un-stimulated, 3 defined by lethargy, hunched posture, and subdued behavior even when stimulated, and 4 defined by bleeding, unresponsiveness, severe weakness, or inability to walk. All mice scoring a 4 were considered moribund and were euthanized based on IACUC-approved criteria. Daily observations were increased to a minimum of twice daily while mice were exhibiting a clinical score of 3.

### Authentic viruses and neutralization assays

The authentic CCHFV isolate CCHFV-IbAr10200 (Genbank Accession 005300) was used in this study. Neutralization assays were conducted as described previously[16, 28]. In brief, heat-inactivated serum was diluted 1:10, and then serial 3-fold dilutions were generated. CCHFV-IbAr10200 was incubated with dilutions for 1 hour at 37 °C. The serum-virus mixture was added to monolayers of VeroE6 cells in a 96-well plate at a final multiplicity of infection (MOI) of 0.08 and incubated for 1 hour at 37 °C. Infection medium was then removed, and fresh cell culture medium without sample was added. 48 h post-infection, culture medium was removed, and plates were fully submerged in 10% buffered formalin and fixed for at least 4 h at room temperature. Plates were removed from formalin and permeabilized with 0.2% Triton-X for 10 min at room temperature and treated with blocking buffer (Cell Stain Buffer; ThermoFisher). Infected cells were detected by consecutive incubation with CCHFV-specific antibody 9D5 (3 µg/ml; BEI NR-40270) and secondary detection antibody (goat anti-mouse) conjugated to Alexa Fluor 488 (1:2000 dilution; Invitrogen). Percent infection was determined using the Cytation5 high-content imaging instrument and data analysis was performed using the Gen5.11 software (BioTek). Percent infectivity values were calculated for each serum sample relative to sera-free, infected wells and normalized to naïve control samples. AUC values were computed for each neutralization curve (baseline Y=0, Ignore peaks <10% of the distance from min to max Y) and log transformed. AUCs were compared by 2-way ANOVA with Turkey corrections for multiple comparisons (GraphPad Prism 10.0.3).

